# Calcium Signalling in Glioblastoma Networks of Different Topologies and Possible Treatments

**DOI:** 10.1101/2025.08.19.671177

**Authors:** Alexandra Shyntar, Thomas Hillen

**Author notes:** Dedicated in memory of Dr. Siv Sivaloganathan, a visionary in mathematical medicine.

## Abstract

Glioblastoma cells form connected cell networks, utilizing tumor microtubes to transmit calcium between cells. A new cell type called “periodic cell” is integral in sustaining calcium signalling in a glioblastoma network. Periodic cells are rare, can sustain consistent intracellular calcium transients, are likely to have KCa3.1 pumps, and have on average more tumor microtubes than other glioma cells. Here we adapt an ordinary differential equation model for intracellular as well as intercellular calcium signalling and apply it to a large glioma cell network. Using the model, three main hypotheses for the driving mechanism of periodic cells were tested: 1. a fixed and elevated IP_3_ concentration, 2. added benefit from influx of calcium due to KCa3.1 pumps, or 3. oscillation in calcium influx into the cell through the plasma membrane. All three hypotheses yield similar calcium oscillation patterns resembling the trends seen in the data of Hausmann et al. 2023. In vivo, glioma networks were shown to have small-world and scale free network properties. We apply our model to small-world, scale-free and random networks. For these networks, we test how communication is inhibited through removal of cells, removal of tumor microtubes, and inhibition of KCa3.1 pumps. All three network types were more vulnerable to random cell damage than to random TM damage. We find that inhibition of KCa3.1 pumps can have a significant impact on the inhibition of network communication, however, to fully degrade the calcium signalling network, all periodic cells must be eradicated, confirming experimental observations.

## 1 Introduction

Glioblastoma is a disease that is highly invasive and resistant to conventional therapies. It has recently been shown that glioblastoma cells can form a communication network using tumor microtubes (TMs) [17]. Tumor microtubes are thin long protrusions that extend from a cells body and connect to other glioma cells through connexin-43 gap junctions [32]. Through these connections, calcium can be transmitted from one cell to another thereby creating calcium transients in the glioblastoma cell cytoplasm and facilitating communication between cells [32, 17]. The cell network in glioblastoma is a property of the tumor bulk, where cell signalling is absent from the invasive part of the tumor [17].

Here, we develop a mathematical model to describe how calcium signalling is transported in glioma networks and to understand how different treatments can inhibit such a communication. To do this, we take the Höfer [20] model for intracellular and intercellular calcium signalling in cells and extend it by testing various hypotheses. The hypotheses pertain to the different ways calcium oscillations may arise in periodic cells. We apply the models to three different networks: small-world, scale-free, and random networks. Then, we test different treatments either a random or a targeted treatment. In a random treatment the removal of cells or TMs is random. In a targeted treatment, the removal focuses on removal of key cells or inhibition of the KCa3.1 pumps. Our model is able to reproduce the experimental data from [17] and it confirms the observation that the key to stop network communication is the inhibition of periodic cells.

### 1.1 Periodic cells and glioblastoma networks

In a recent paper, Hausmann et al. [17] performed a series of mice experiments by implementing S24 glioblastoma strands into mice and observing the calcium transients. The experiments of Hausmann et al. [17] revealed the existence of a subpopulation of cells in glioblastoma, called periodic cells, which make up less than 7% of the total glioblastoma cell population. Periodic cells are more involved in cell-to-cell signalling as they produce the highest amount of calcium transients and produce periodic calcium oscillations [17]. Further, periodic cells generally have more TMs than other cells and they help increase cell proliferation as well as decrease cell death of nearby cells [17]. Calcium activity tends to be less in glioblastoma cells that are further away from periodic glioblastoma cells, as periodic cells are the ones that trigger calcium activity in non periodic cells [17]. Hausmann et al. [17] found that the number of periodic cells in a network can change over time. In particular, ATP can increase the amount of periodic cells. Further, after some damage to the cell network new periodic cells can appear [17]. Moreover, periodic cells tend to express KCa3.1 more than non periodic cells [17]. KCa3.1 is a potassium channel that effluxes potassium ions out of the cell, which promotes the entry of calcium ions into the cell [17]. KCa3.1 expression has been previously linked to poorer patient outcome, resistance to treatments such as temozolomide chemotherapy, and to glioma motility [17].

Upon further analysis of the glioblastoma cell network, Hausmann et al. [17] discovered that the network has both small world and scale-free network properties. In the scale-free networks, about 5% of glioblastoma cells are highly connected to other glioblastoma cells, and thus these cells act as functional network hubs. The cells acting as functional network hubs often have periodic activity as well. Hausmann et al. [17] showed that targeting and eliminating these network hubs leads to the degradation of the network, which gives rise to promising treatments for glioblastoma.

### 1.2 Treatments

In this subsection, we discuss the treatments that Hausmann et al. [17] performed. One treatment consisted of using a laser to either remove glioblastoma cells randomly or specifically. Randomly killing cells in the network did not significantly disrupt the cell network as networks that have network hubs are highly resistant to random damage [17]. The removal of periodic cells lead to the fragmentation of the network, which significantly decreased the networks functionality [17]. Moreover, Hausmann et al. [17] found that removing periodic cells also led to higher cell death of the glioblastoma cells that were connected to the network.

Hausmann et al. [17] used TRAM-34 and senicapoc to inhibit the KCa3.1 channel. Inhibiting the KCa3.1 channel reduced the calcium activity, the calcium frequency of periodic cells, glioblastoma cell proliferation, as well as tumor invasion [17]. Hausmann et al. [17] further experimented with genetic knockout (via CRISPR) and knockdown (via short hairpin RNA (shRNA)) of KCa3.1. They found that periodic activity as well as the proliferation rate of glioblastoma cells decreased post genetic knockout or knockdown of KCa3.1. This inhibition of KCa3.1 did not affect irregular non-periodic calcium activity and proliferation of tumors that do not form cell networks [17]. Hausmann et al. [17] found that the network could be repaired after the inhibition of KCa3.1 by introducing KCa3.1-proficient wild-type cells. The KCa3.1-proficient wild-type cells produced more periodic cells over time hence the cell network got restored and the proliferation rate of glioblastoma restored to the original one pre treatment.

Other treatments that were tested by Hausmann et al. [17] involved decreasing calcium entry into the cell (e.g. via calcium chelation), gap junction or TM inhibition, and ATP inhibition. Hausmann et al. [17] found that calcium chelation near periodic cells significantly reduced the proliferation rate of nearby cells. The further away the glioblastoma cell was from the periodic cell the less impact calcium chelation had on its proliferation rate. Hausmann et al. [17] found that inhibiting TM formation or gap junctions, significantly reduced the amount of intercellular calcium activity as well as the proliferation of glioblastoma cells. Furthermore, Hausmann et al. [17] found that inhibiting extracellular ATP transfer did not significantly impact the calcium activity within the cellular network.

### 1.3 Calcium mechanisms within the vell

In this section, we outline the primary mechanisms involved with the calcium dynamics in a glioblastoma cell, which were noted by Hausmann et al. [17]. We begin with calcium entry into the cell. Calcium ions can enter through the store-operated calcium channels (SOCCs), receptor-operated channels, or voltage-gated channels (VGCs) [28]. Hausmann et al. [17] emphasized the Orai channel for calcium entry, which falls into the SOCC category. SOCCs are triggered to open by proteins such as the stromal interaction molecules (STIM), which are released when calcium in the endoplasmic reticulum (ER) gets depleted [13]. Therefore, as the calcium gets depleted from the ER, the Orai channels open. We illustrate calcium entry through these pumps as a black arrow crossing the plasma membrane (green curve) in Figure 1. Additionally, calcium can be transmitted through the gap junctions from one cell to another to increase the calcium concentration in the cell cytoplasm (this is not illustrated in Figure 1). The concentration of calcium in the cell cytoplasm can also be increased by releasing calcium from the ER. For this, Phospholipase C (PLC) must convert phosphatidylinositol-4,5-bisphosphate into inositol-1,4,5-triphosphate (IP_3_) [27]. Then IP_3_ can bind to the IP_3_ receptor (IP_3_R) to open it in order to release calcium [27] (see Figure 1). The activity of IP_3_R with respect to the calcium concentration in the cell cytoplasm can be described by a bell shaped function where IP_3_R opens at low concentrations of calcium in the cell cytoplasm (positive feedback from calcium) and closes at high concentrations of calcium in the cell cytoplasm (negative feedback from calcium) [2]. In Figure 1, this is illustrated by the blue arrow starting from the calcium ions and pointing to the IP_3_R.

**Figure 1:**
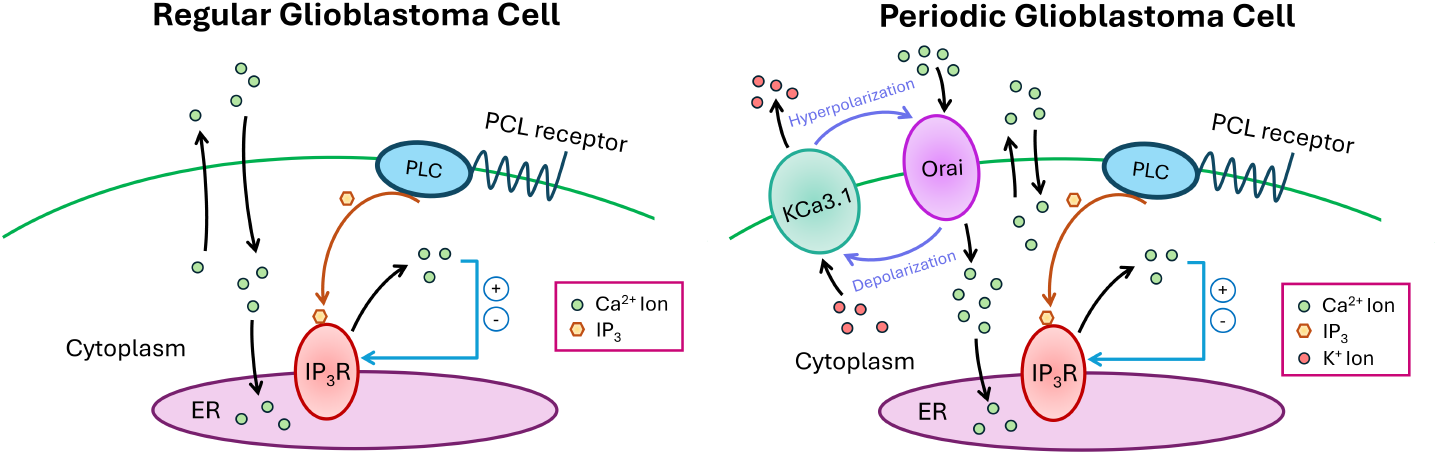
Illustration of the calcium dynamics within a regular glioblastoma cell and a periodic glioblastoma cell. Images are adapted from Figure 4a in [17].

The calcium ions can also leave the cell cytoplasm by either leaving through the cell membrane or entering the ER. The calcium ions can leave the plasma membrane via plasma membrane Ca^2+^ ATPase (PMCA) pump for example [3], or can enter the ER through the sarcoplasmic/endoplasmic reticulum calcium ATPase (SERCA) pump [8]. This is indicated by black arrows in Figure 1.

The key difference between regular glioblastoma cells and periodic glioblastoma cells is that periodic glioblastoma cells have an additional potassium channel: KCa3.1. The KCa3.1 channel is a calcium activated channel and is independent of voltage [35]. When the potassium ions efflux out from the KCa3.1 hyperpolarization of the plasma membrane occurs. Hyperpolarization helps more calcium ions to pass through due to the negative membrane potential. The plasma membrane depolarizes which can activate the KCa3.1 channels again thereby creating a positive feedback loop [17]. This is illustrated in Figure 1 in the right image. In addition, the KCa3.1 can be activated by the P2Y_2_ receptor which is a PLC receptor [17].

### 1.4 Previous mathematical modelling approaches for calcium transients in cells

There have been several approaches in the literature to model the intracellular calcium oscillations, where it was shown that calcium oscillations can stem from the interplay between the calcium in the cytoplasm and ER or the oscillation in the IP_3_ concentration. Meyer et al. [29] proposed an ODE system that accounted for both the calcium concentration in the cytoplasm and the ER as well as the variable concentration of IP_3_. The Meyer et al. [29] model showed that oscillations stem from the varying IP_3_ concentration. An early model of Kuba et al. [24] proposed an ODE system for calcium oscillation in ganglion cells without incorporation of IP_3_. Later, Kuba et al. [24] model was extended by Goldbeter et al. [16]. The new model accounted for IP_3_ and assumed that there are two calcium stores inside the cell that release calcium into the cell’s cytoplasm. One calcium store is sensitive to IP_3_ and the other is sensitive to calcium concentration. The model presented by Goldbeter et al. showed that one way the calcium oscillations can be produced is through a sufficiently large IP_3_ concentration. Unlike the model presented by Meyer et al. [29], IP_3_ oscillation is not necessary to produce calcium oscillations in the cytoplasm. Dupont et al. [11] showed that the model proposed by Goldbeter et al. [16] can reproduce the calcium oscillations with sufficiently large IP_3_ concentration by dropping the assumption that there are two different calcium stores inside the cell. Instead, there is one calcium store (the ER) that is sensitive to both the cytoplasmic calcium concentration and the IP_3_ concentration.

Following the experiments of Bezprozvanny et al. [2], which studied the IP_3_ channel in more detail, more calcium models were proposed that incorporated their findings. Bezprozvanny et al. [2] showed that the channels opening probability resembles a bell curve, where it opens at low calcium concentrations and closes at high calcium concentration in the cytoplasm while additionally requiring the activation from IP_3_. De Young et al. [10] proposed a model similar to Goldbeter et al. [16] with a dynamic IP_3_ concentration while incorporating the findings of [2]. That is, De Young et al. [10] model incorporates the three main components of an IP_3_ channel which are: regulation by IP_3_, activation by calcium, and inactivation by calcium. Later, Li et al. [26] simplified the model of De Young et al. [10] by exploiting time scales in the rates of the three main components. Later, Höfer [20] proposed a calcium model for calcium oscillations in hepatocytes while incorporating calcium signalling from other hepatocytes via gap junctions. Höfer [20] used the simplification of Li et al. [26] to model the release of calcium from the ER and proposed a model similar to Dupont et al. [11], meaning that the IP_3_ concentration can remain fixed and oscillation of calcium in the cell cytoplasm can be observed in the simulations. Here, we extend the Höfer [20] model to apply for glioblastoma cells, and discuss the Höfer [20] model in more detail in one of the following sections.

### 1.5 Mathematical models for glioma

There have been numerous mathematical models for modelling glioma with various approaches. To study cell structure, adhesion, and cell motility, agent-based models such the cellular Potts model have been used [14]. Also, reaction-diffusion models are a popular choice when modelling glioma growth and spread [39, 25, 21]. The reaction-diffusion equations have been extended to transport equations with anisotropic diffusion, allowing to easily incorporate the available diffusion tensor imaging (DTI) data for cell movement along white matter tracts [15, 33, 7, 38]. Mechanistic models for glioma growth and spread have also been proposed to study effects of nutrient availability [6] or tissue deformation in addition to cancer growth and spread [34].

The glioma models discussed above do not account for the recent discovery of tumor microtubes (TMs) that help with cell proliferation, calcium signalling, and cancer invasion [32, 17]. Recently, Hillen et al. [19] proposed a transport equation model accounting for the TMs ability to invade tissues as well as glioma nuclei migration using TMs, in addition to accounting for cell proliferation and movement. Shyntar et al. [36], proposed a simpler partial differential equation model accounting for the TMs ability to aid with cell proliferation and directed motion. The model proposed by [36] specifically focused on the case of glioma regrowth and spread post surgery in the mice experiments performed by [41].

Our focus here is on calcium signalling in glioma and its transmission in a glioma network. Here we ignore cell division and movement, as these involve much longer time scales. For model modification, we perform a similar but simpler approach taken from Catacuzzeno et al. [4], who extended the calcium model proposed by Höfer [20]. The model of Catacuzzeno et al. [4, 5] extends the model proposed by Höfer [20] by incorporating the effect of KCa3.1 and the resulting transmembrane potential effect on calcium influx into the cell. Catacuzzeno et al. [4] account for how the calcium inflow, potassium outflow, and general ion leakage effects the transmembrane potential. They find that an oscillating calcium influx into the cell, in contrast to a constant influx as used by Höfer [20], increases the amplitude, the duration, and the frequency, of the calcium spikes [4, 5]. Furthermore, Catacuzzeno et al. [4] included voltage dependence for calcium transmission through gap junctions to other cells, which was not done in the model proposed by Höfer [20]. Like the results in Höfer [20], Catacuzzeno et al. [4] showed that their model is able to show calcium spike synchronicities between different cells.

### 1.6 Outline of the paper

In Section 2 we formulate our main model, which is based on the calcium model of Höfer [20]. It is a model that balances calcium fluxes between the cell cytoplasm, the endoplasmic reticulum (ER) and the outside of the cell. We recall the basic properties of the Höfer model in Section 3 and A.2, where we present its bifurcation structure. Depending on the parameters, solutions either converge to a steady state, or they converge to a periodic orbit. We show that the non-periodic cells form an excitable system, where a sufficiently large stimulation can trigger a full calcium spike. Periodic cells, however, live in the Hopf-region, where sustained oscillations are possible. Section 4 is used to identify model parameters and to calibrate the model on the data of Hausman [17]. In Section 5 we go beyond the experiments of Hausmann and analyse large glioma networks that show scale free and small world properties. We compare those with purely random networks. In Section 6 we add treatments of the glioma networks that are intended to disrupt the calcium wave transmission. We find that random removal of cells or random removal of tumor microtubes has limited impact, while inhibition of KCa3.1 pumps and removal of periodic cells is a viable treatment option, confirming the experimental observations. We close with a Conclusion Section 7.

## 2 Model formulation

### 2.1 Höfer (1999) model

Previously, Höfer [20] proposed a mathematical model for intercellular calcium oscillations in hepatocytes. Here, we will discuss the model in more detail and after that, we will use it for calcium signalling in glioblastoma cells. The model proposed by Höfer [20] models the five essential calcium fluxes in a cell: the calcium flux coming into the cell through the cell plasma membrane (*J*_*in*_), the calcium flux coming out of the cell through the cell plasma membrane (*J*_*out*_), the calcium flux coming out of the ER into the cell cytoplasm (*J*_*rel*_), the calcium flux going into the ER from the cytoplasm (*J*_*SERCA*_), and the gap-junctional influx of calcium coming from other cells (*J*_*G*_) (see Figure 2 for summary). Note that *J*_*G*_ is also used to represent the efflux of calcium from a given cell to other cells, as we will discuss later.

**Figure 2:**
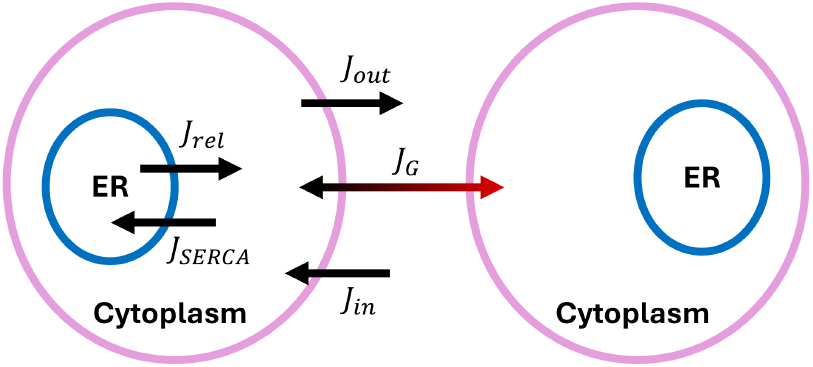
Illustration of the cell fluxes of the Höfer model [20]. The flux *J*_*G*_ may be bidirectional with or without preference, or be unidirectional depending on how it is defined in the model.

Let *x* be the concentration of calcium in the cell cytoplasm and *y* be the concentration of calcium in the cell ER. The general mathematical model proposed by Höfer [20] for two cells illustrated in Figure 2 incorporating calcium buffer dynamics within the cytoplasm and the ER is given by

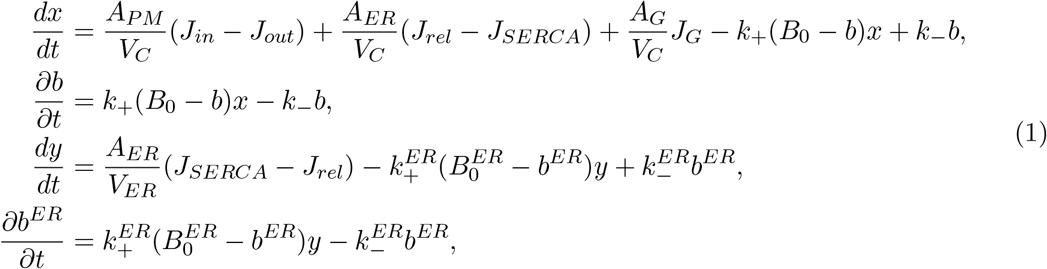

where *A*_*P M*_ is the area of the cell plasma membrane, *A*_*ER*_ is the area of the ER membrane, *A*_*G*_ is the area of the gap junctional connection, and *V*_*C*_ and *V*_*ER*_ are the cytosolic volumes of the cytoplasm and ER respectively. The last two terms in each equation in (1) account for the buffering of calcium where *B*_0_ is the total concentration of the buffer (bound and unbound) in cytoplasm, *b* is the concentration of the buffer bound to calcium, *k*_+_ is the rate at which the buffer binds to calcium, and *k*_*−*_ is the rate at which the buffer unbinds from the calcium in the cytoplasm. Note that *B*_0_ = *b* + *B*_*u*_ where *B*_*u*_ is the unbound buffer. The parameters with the superscripts in the last two terms of the third equation and in the fourth equation in (1) have the same meanings as the ones without the superscripts except that the parameters with the subscripts apply to the concentrations and rates in the ER. Similarly, 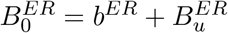.

Under quasi-steady state assumptions for *b* and *b*^*ER*^, the above model (1) can be rewritten as (see Appendix A.1 for details)

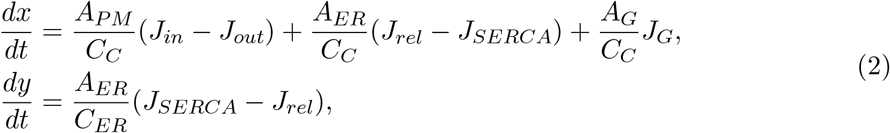

where *C*_*C*_ and *C*_*ER*_ are the effective cytosolic volumes or calcium capacity in the cytoplasm and ER, respectively.

Using the transformation

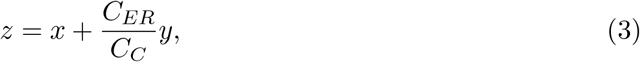

we can rewrite (2) as

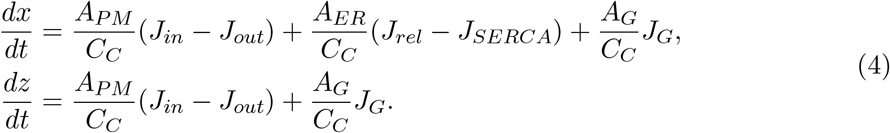

Now we go over the specific forms of the flux terms used by Höfer [20]. By assuming that the calcium influx increases with the amount of IP_3_ (*P*) up to a saturation point, the specific form of *J*_*in*_ used by Höfer [20] is

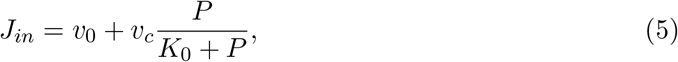

where *v*_0_ is the background leakage of calcium through the plasma membrane. The *v*_*c*_-term describes the influx of calcium through SOCC pumps, that results from depletion of calcium from the ER, that was activated with IP_3_ [20]. Here, *K*_0_ is the half-saturation constant.

The specific form of *J*_*out*_ used in [20] is

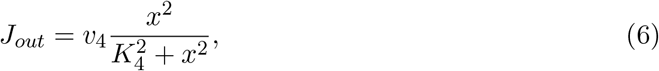

where *v*_4_ is the maximum outflow of calcium activity through the plasma membrane and *K*_4_ is the half-saturation constant.

The *J*_*rel*_ form used by Höfer [20] comes from the simplification of Li et al. [26] and is given by

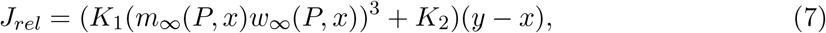

where *K*_1_ is the maximal IP_3_R-mediated release rate and *K*_2_ is the small leak flux. The function *m*_*∞*_(*P, x*) models the faster process of channel opening via IP_3_ in comparison to calcium activation/deactivation of the channel while being independent from the kinetics of these slower processes [26]. Its explicit form is given by

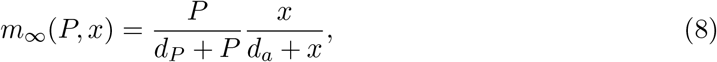

where *d*_*P*_ and *d*_*a*_ are half-saturation constants. The function *w*_*∞*_(*P, x*) models the slowest process, which is the channel inactivation via calcium and the function depends, the kinetics of the faster processes [26]. Höfer [20] assumes that no significant role is played by the timing course of IP_3_R inactivation on the timing of the calcium spike. Because of this assumption, the explicit form of *w*_*∞*_(*P, x*) used in [20] is given by

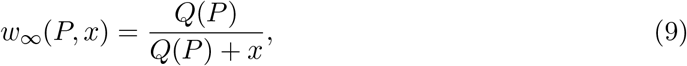

and

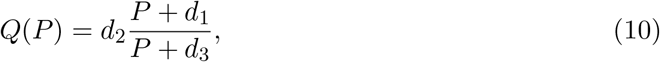

where *d*_1_, *d*_2_, *d*_3_ are reaction rate constants for the binding processes.

The explicit form for *J*_*SERCA*_ used in [20] is given by

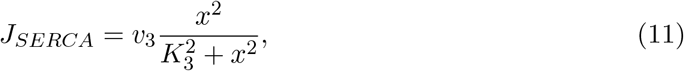

where *v*_2_ is the maximum inflow activity through the ER membrane and *K*_2_ is the half-saturation constant.

The explicit form of *J*_*G*_ used in [20] is given by

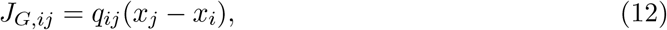

where *q*_*ij*_ is the gap junctional permeability, *x*_*i*_ and *x*_*j*_ refer to calcium cytoplasm concentration of cells *i* and *j*, respectively.

Setting 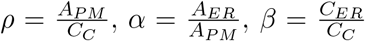, and 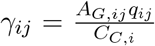, we can rewrite the model (4) for many cells *x*_*i*_ *i* = 1, …, *n* that are coupled via the gap junctions with permeability rate *γ*_*ij*_ as follows. For each *i* = 1, …, *n*

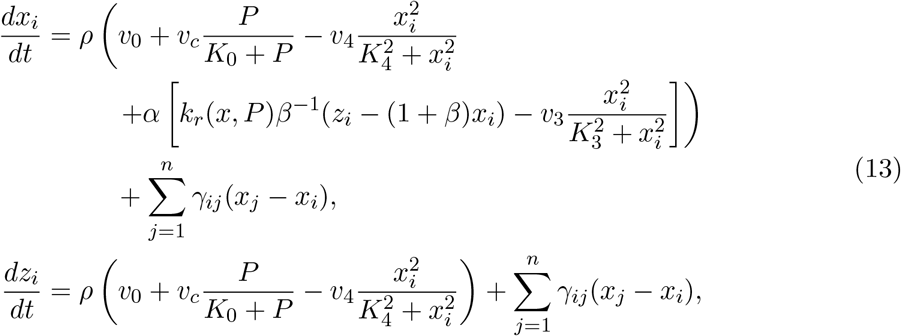

with

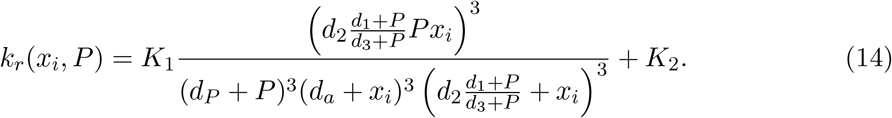

We abbreviate

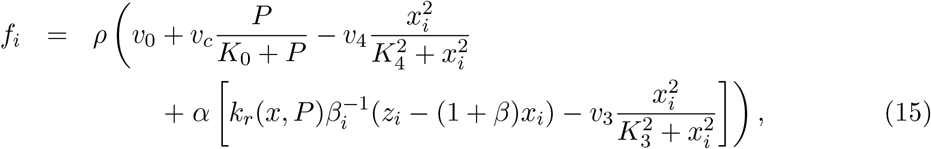

and

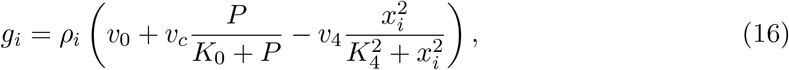

which allows us to rewrite model (13) as

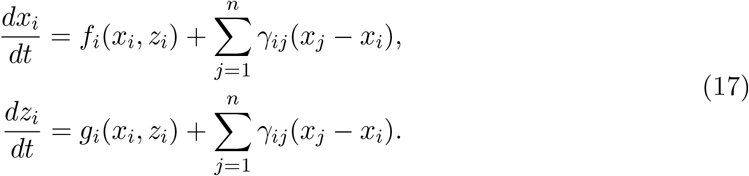

where *z*_*i*_ is given by (3).

Given that there are *n* glioblastoma cells, we set *m* (*m < n*) of them to be periodic cells. Hence, our glioblastoma cell population splits into two distinct categories: *m* periodic cells that have periodic calcium transients when unperturbed and *n* – *m* glioblastoma cells that have their cytoplasmic calcium settle into equilibrium over time. The difference between periodic and non-periodic cells in (17) relates to different forms of the activation functions *f*_*i*_ and *g*_*i*_. The functions defined in (15) and (16) relate to the non-periodic glioma cells. The activation functions for periodic cells have slightly modified parameters and possibly a modified input, and we define those later in (20) and (21), respectively.

## 3 Analysis of Höfer’s model

In this section, we summarize the behavior of the model (4) for one cell. The detailed mathematical analysis can be found in A.2. For a single cell, in order to have intracellular calcium oscillations, the model (4) must exhibit periodic solutions. Indeed, the model exhibits two Hopf bifurcations, a forward Hopf for *P* = 1.45 and a backward Hopf for *P* = 8.892. For *P* ∈ (1.45, 8.892) we have stable periodic orbits. As we use *P* as a bifurcation parameter, all other parameters are held constant (see Table 1). The phase portraits of the model (4) in Figure 3 illustrate this bifurcation structure.

**Table 1:** Summary of the parameters used. The tuple in the last column denotes the references for the parameters. The first entry and second entries of the tuple are the references to the parameters in the Periodic Cell column and the Non-Periodic Cell column respectively. If the reference says estimated, then we chose these values as they best matched the data. The parameters with multiple values correspond to different values taken under different hypotheses. For *P*, the first value, the second value, the third value correspond to the values taken under hypothesis H1, H2 and H3, respectively. Note that 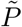 applies only to periodic cells. Similarly for *κ*, its value is relevant only under H2.

| Parameter | Periodic Cell | Non-Periodic Cell | Units | Reference |
| --- | --- | --- | --- | --- |
| $\rho$ | 0.011 | 0.011 | $1/\mu\text{m}$ | estimated |
| $\beta$ | 0.1 | 0.1 | - | ([20],[20]) |
| $\alpha$ | 2 | 2 | - | ([20],[20]) |
| $v_4$ | 3.6 | 3.6 | $(\mu\text{M } \mu\text{m})/\text{s}$ | ([20],[20]) |
| $K_4$ | 0.12 | 0.12 | $\mu\text{M}$ | ([20],[20]) |
| $v_3$ | 9 | 9 | $(\mu\text{M } \mu\text{m})/\text{s}$ | ([20],[20]) |
| $K_3$ | 0.12 | 0.12 | $\mu\text{M}$ | ([20],[20]) |
| $K_1$ | 40.0 | 40.0 | $\mu\text{m/s}$ | ([20],[20]) |
| $\tilde{P}, P, P$ | 1.8, 1.4, 1.4 | 1.4, 1.4, 1.4 | $\mu\text{M}$ | estimated |
| $d_P$ | 0.2 | 0.2 | - | ([20],[20]) |
| $d_a$ | 0.4 | 0.4 | $\mu\text{M}$ | ([20],[20]) |
| $d_2$ | 0.4 | 0.4 | $\mu\text{M}$ | ([20],[20]) |
| $d_1$ | 0.3 | 0.3 | $\mu\text{M}$ | ([20],[20]) |
| $d_3$ | 0.2 | 0.2 | $\mu\text{M}$ | ([20],[20]) |
| $K_2$ | 0.02 | 0.02 | $\mu\text{m/s}$ | ([20],[20]) |
| $v_0$ | 0.2 | 0.2 | $(\mu\text{M } \mu\text{m})/\text{s}$ | ([20],[20]) |
| $v_c$ | 4.0 | 4.0 | $(\mu\text{M } \mu\text{m})/\text{s}$ | ([20],[20]) |
| $K_0$ | 4 | 4 | $\mu\text{M}$ | ([20],[20]) |
| $\gamma$ | 0.0084 | 0.0084 | $1/\text{s}$ | estimated |
| $\kappa$ | -, 1.3, - | -, 1, - | - | - |

**Figure 3:**
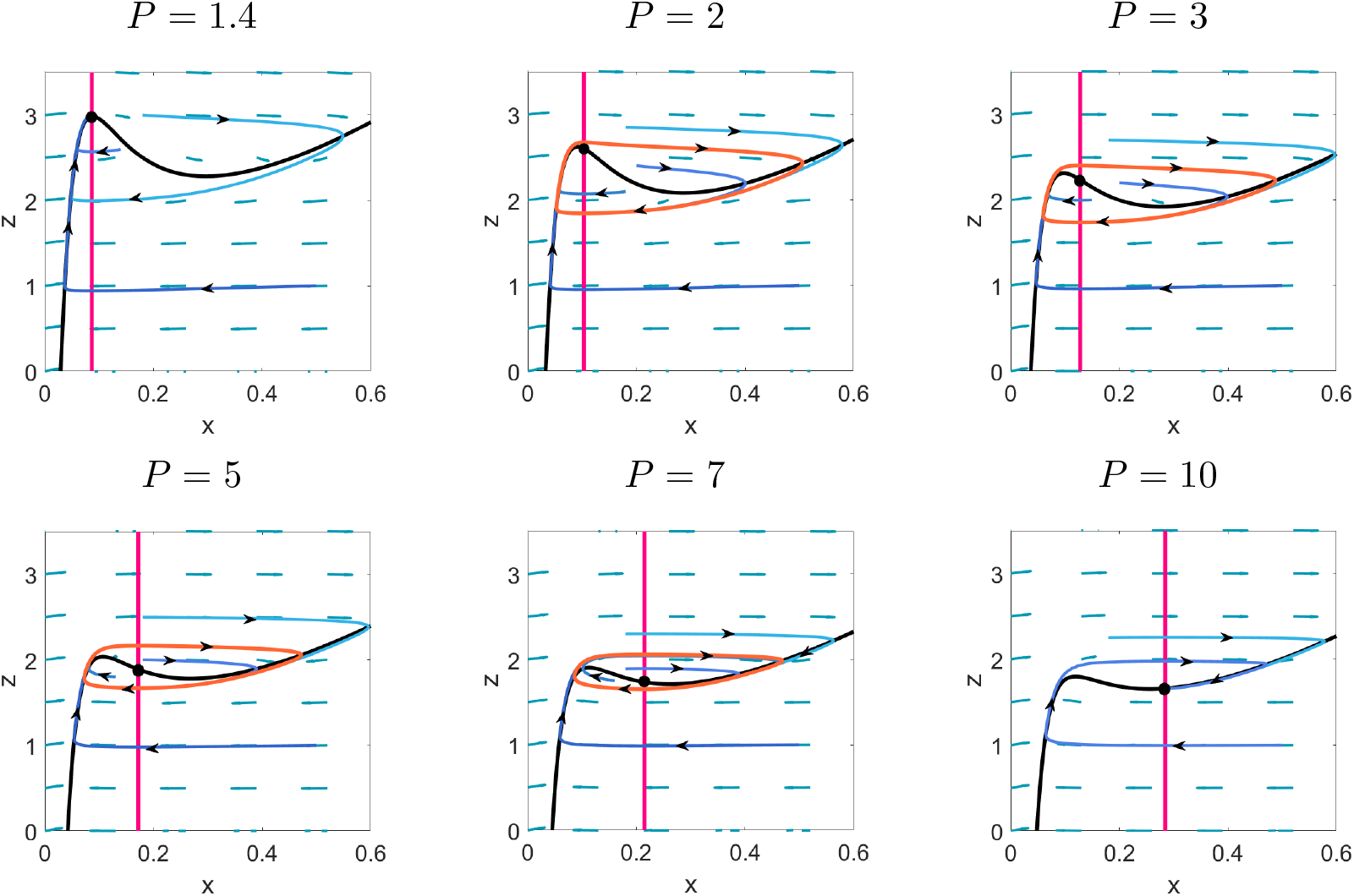
Phase portraits with parameters used from Table 1 with varying *P*. The value of *P* is denoted above each figure. The pink line is the *z* nullcline and the black curve is the *x* nullcline. The blue curves are example trajectories and the periodic orbit is shown in orange. The parameters used are given in Table 1.

We next study the case of two cells, where a periodic cell signals to a non periodic cell. To do this we set *P* = 1.8 for a periodic cell and *P* = 1.4 for a non-periodic cell. All other parameters are equal between cells and can be found in Table 1. Hence, the model is

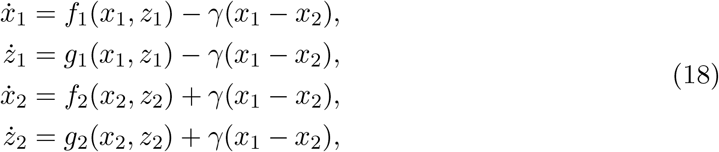

where *x*_1_ and *z*_1_ correspond to the calcium concentrations in the periodic cell and *x*_2_ and *z*_2_ correspond to the calcium concentrations in the non-periodic cell.

The resulting solutions and phase portraits are shown in Figure 4. From the phase portraits, we see that the solution trajectory of a receiver cell requires a stimulus large enough to push it across the *x* nullcline and *z* nullcline in order to have a pronounced spike in the *x* solution. In row 4, column 2 of Figure 4, we see an example of where the impulse was not enough to send the solution trajectory on an excursion and yield a large spike as seen in the other images in row 4. This shows that the concentration of the cytosolic and ER calcium needs to be sufficiently close to the equilibrium, in order for the calcium concentration coming from a signalling cell to be enough to yield a calcium spike in the receiver cell. Biologically there are two interpretations of this phenomenon. Either the calcium concentration in the ER needs to be sufficiently large to release calcium into the cytoplasm after receiving calcium or the increase in the cytosolic calcium after receiving calcium from the signalling cell inhibited the IP_3_ receptors from releasing calcium from the ER.

**Figure 4:**
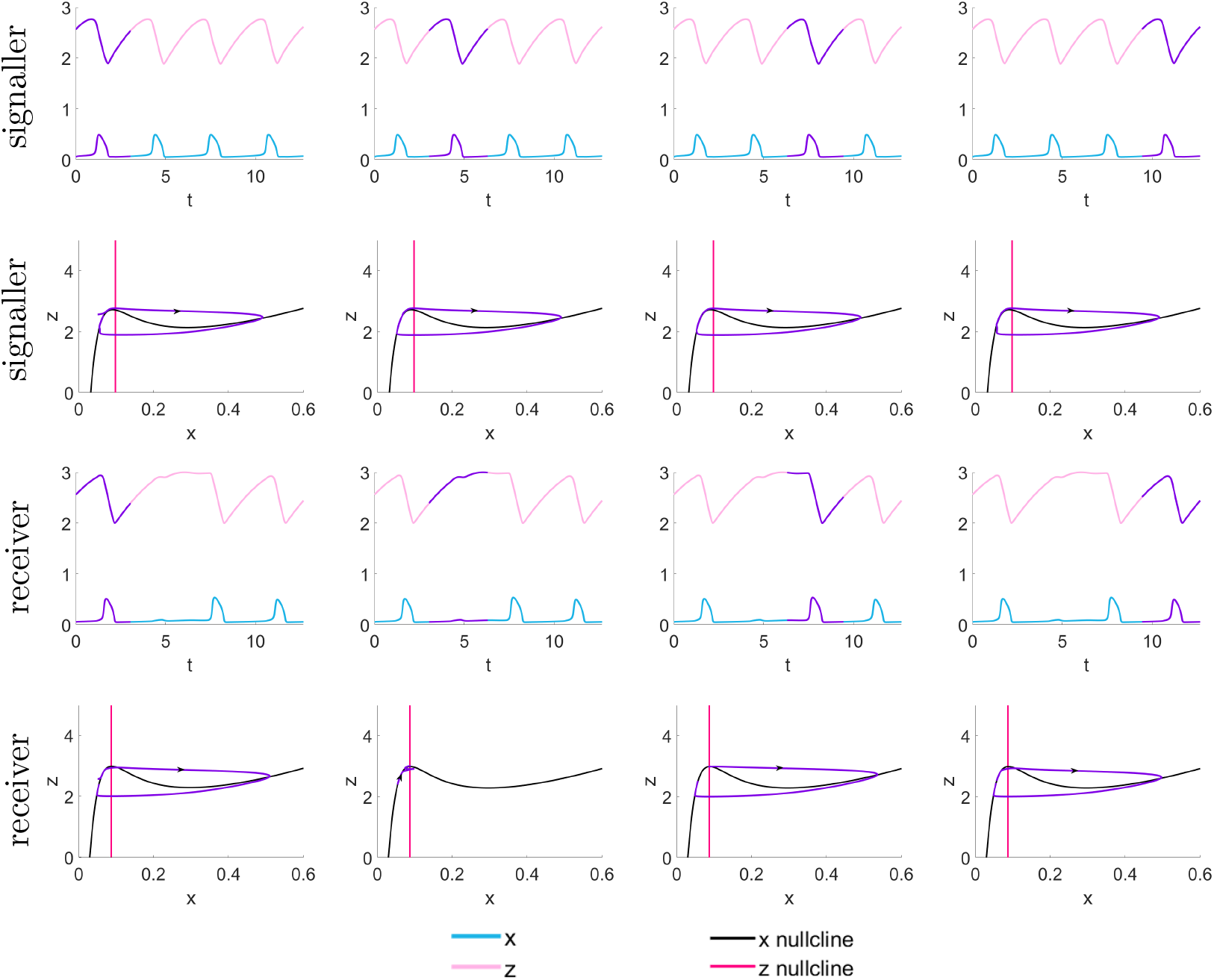
Solutions and phase portraits from system (18) where *P* = 1.8 in signaller cells and *P* = 1.4 receiver cells. Time *t* has a unit of minutes. All other parameters are equal and found in Table 1. In row 1, the resulting concentrations *x*_1_ and *z*_1_ are shown, where blue is the *x*_1_ concentration and pink in the *z*_1_ concentration. The purple highlights the solution part that is illustrated in the phase portrait in row 2. For the phase portraits in row 2, *x*_1_ is on the horizontal axis and *z*_1_ is on the vertical axis. The black curve is the *x*_1_ nullcline and the pink curve is the *z*_1_ nullcline. Similarly, in row 3, the resulting concentrations of *x*_2_ (blue) and *z*_2_ (pink) are shown. Purple highlights the solution part that is illustrated a phase portrait in row 4. The black curve is the *x*_2_ nullcline and the pink curve is the *z*_2_ nullcline.

## 4 Numerical Simulation of the Glioma-Network Model (17)

Here we come back to the network model (17) and consider networks of various sizes and topologies.

### 4.1 Model parameters

Due to a lack of specific parameters for glioma, we set most of the parameters to have the same values as those used by Höfer [20] for hepatocytes. The parameters are summarized in Table 1. The parameters related to calcium dynamics are unchanged but the parameters that are more specific to glioblastoma will be discussed now.

- **Area to volume ratio** *ρ*: We set *ρ* = 0.011*µ*m^*−*1^ (different form the value used in [20]). Based on experimental images [17] the glioblastoma cells have an average approximate radius of 8*µ*m. Assuming that the cell has a spherical shape, we get an approximation of *A*_*P M*_ ≈ 804*µ*m^2^ and the total cell volume of *V* ≈ 2145*µ*m^3^. Cytosol can occupy about 50-64% [31, 42] of the total cell volume, so taking an intermediate value of 57% we get that the volume of cytosol *V*_*C*_ ≈ 1223*µ*m^3^. Now *ρ* = *A*_*P M*_ */*(*V*_*C*_*X*_*cytosol*_) where *X*_*cytosol*_ is the cytosolic buffering capacity (see Appendix equation (32)). The range of possible *X*_*cytosol*_ is [20,100] [20] and we choose *X*_*cytosol*_ ≈ 60. Then,

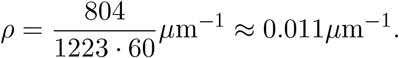
- **Volume ratio** *β*: We keep *β* = *C*_*ER*_*/C*_*C*_ = 0.1 the same as in [20], which we justify as follows. We assume ER makes up about 10% of the cells volume where smooth ER occupies a third of that volume like in a liver cell [20], so *V*_*ER*_ ≈ (0.1*/*3)*V*. As above, *V*_*C*_ ≈ 0.57*V*. ER holds more calcium than cytosol [9] so we assume *X*_*ER*_ = 1.7 *X*_*cytosol*_. So

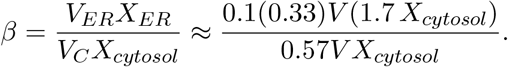
- **Area ratio** *α*: To obtain *α* = 2 we have the same assumption as in Höfer [20] where the smooth ER area is twice as large as the area of the cytoplasm to yield *α* = 2.
- **Gap junction permeability rate** *γ*: The parameter *γ* is variable in Höfer so here we take it to be as a value that best replicates the trends seen in the data.
- **Concentration of IP**_3_ *P* : We choose to set *P* = 1.4 as a base value for non-periodic cells, as this value is outside the range of *P* that triggers calcium oscillations. For periodic cells in hypothesis H1 (see section 4.2), we choose 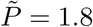 as that makes them sustain intracellular calcium oscillations.

The initial conditions we use are the same as the ones used by Höfer [20]:

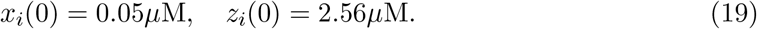

### 4.2 Mechanisms for periodic cells

Due to the complexity of the plasma membrane and their channels, to incorporate the effect of the KCa3.1 we test the following hypotheses H1-H3. Recall, that the activation functions *f*_*i*_ and *g*_*i*_ in (17) were defined as

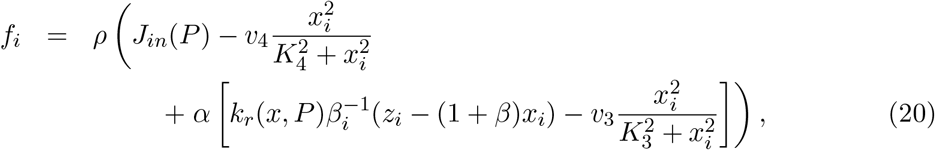

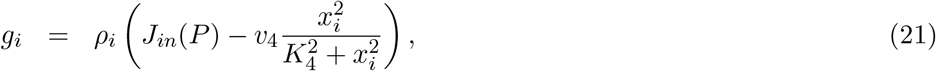

with

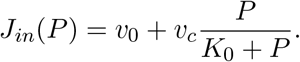

We modify those for periodic cells according to the following cases.

H1. We assume KCa3.1 directly affects *P* in periodic cells. This is motivated by the assumption that the presence of KCa3.1 elevates *P* to a higher value 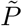 such that the periodic cells are able to have sustained calcium oscillations in model 13. Hence we use 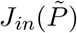 in the expressions for *f*_*i*_ and *g*_*i*_.
H2. We incorporate an indirect impact of KCa3.1 by introducing a sensitivity parameter *κ* into expression of *J*_*in*_, which then becomes

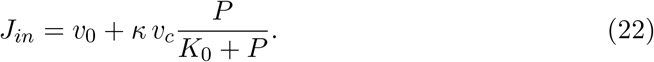

This hypothesis is motivated by the fact that KCa3.1 pumps increase the inflow of calcium into the cell. The *v*_*c*_, as mentioned previously, models the inflow of calcium through SOCC pumps, hence *κ* enhances that flow. We set *κ* to be sufficiently large (*κ* ≥ 1) in order to trigger calcium oscillations in periodic cells under model (13).
H3. KCa3.1 causes a periodic influx into the cell at this particular periodic function

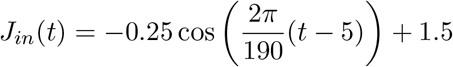

This hypothesis is motivated by the oscillatory *J*_*in*_(*t*) that resulted from the modelling by Catacuzzeno et al. [4]. We simply choose an oscillating function for hypothesis H3 and set its function range based on the range of the influx function used in [4]. The period of the cosine function was taken such that it yields calcium spikes with similar frequency as seen in the data (see Figure 6). For the non-periodic cells, the calcium dynamics are given by model (13) with parameters from Table 1.

**Figure 5:**
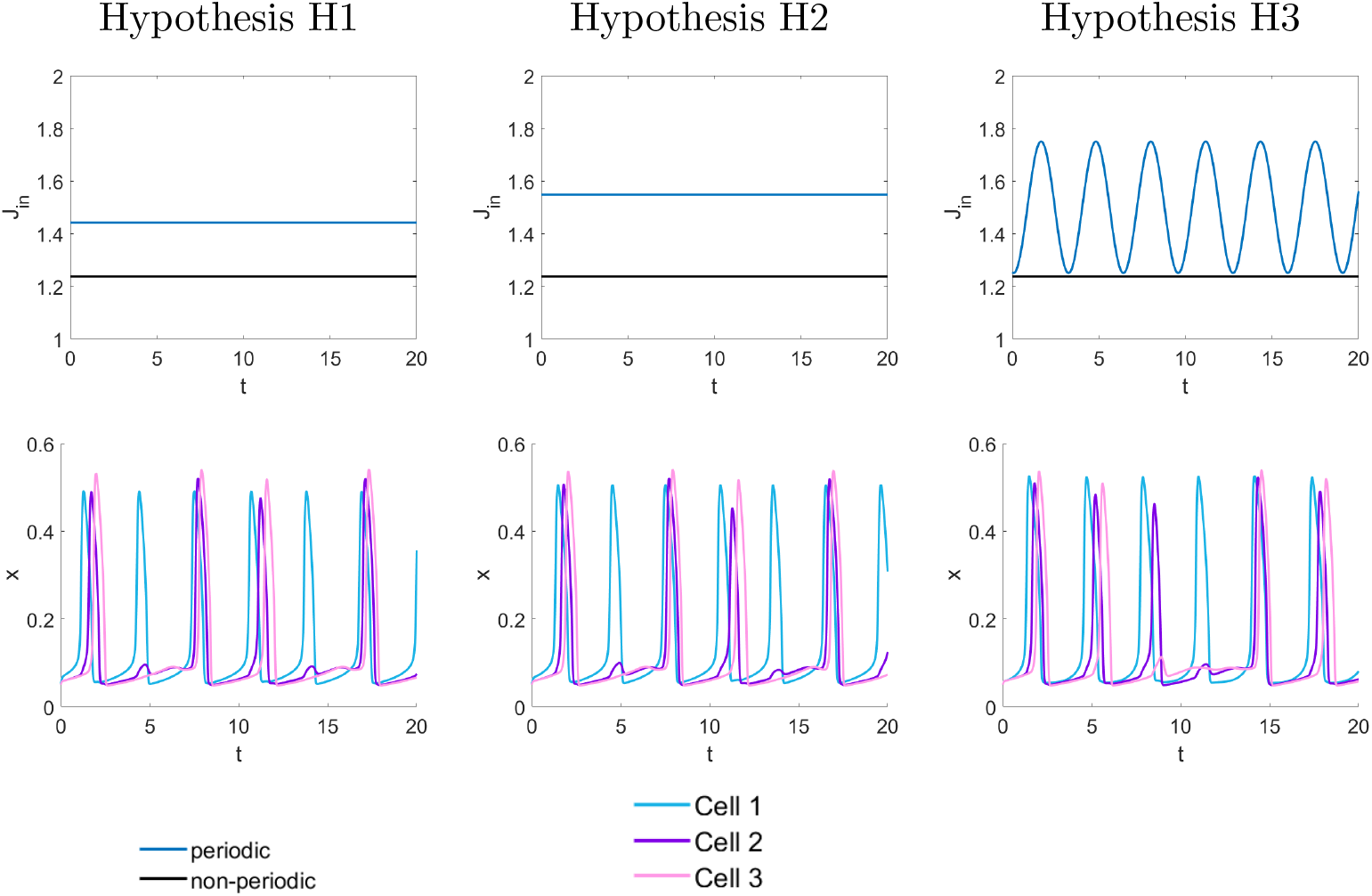
In the first row, the resulting *J*_*in*_ under each hypothesis are illustrated where time has a unit of minutes. In the second row, the solutions of the glioma-network model (17) are shown where parameters are taken from Table 1. Cell 1 is periodic, which signals to cell 2 (non-periodic) and cell 2 signals to cell 3 (non-periodic). Note that in the solutions only the cytosolic calcium concentration, *x*, is plotted and the concentration in the ER, *z*, is not illustrated.

**Figure 6:**
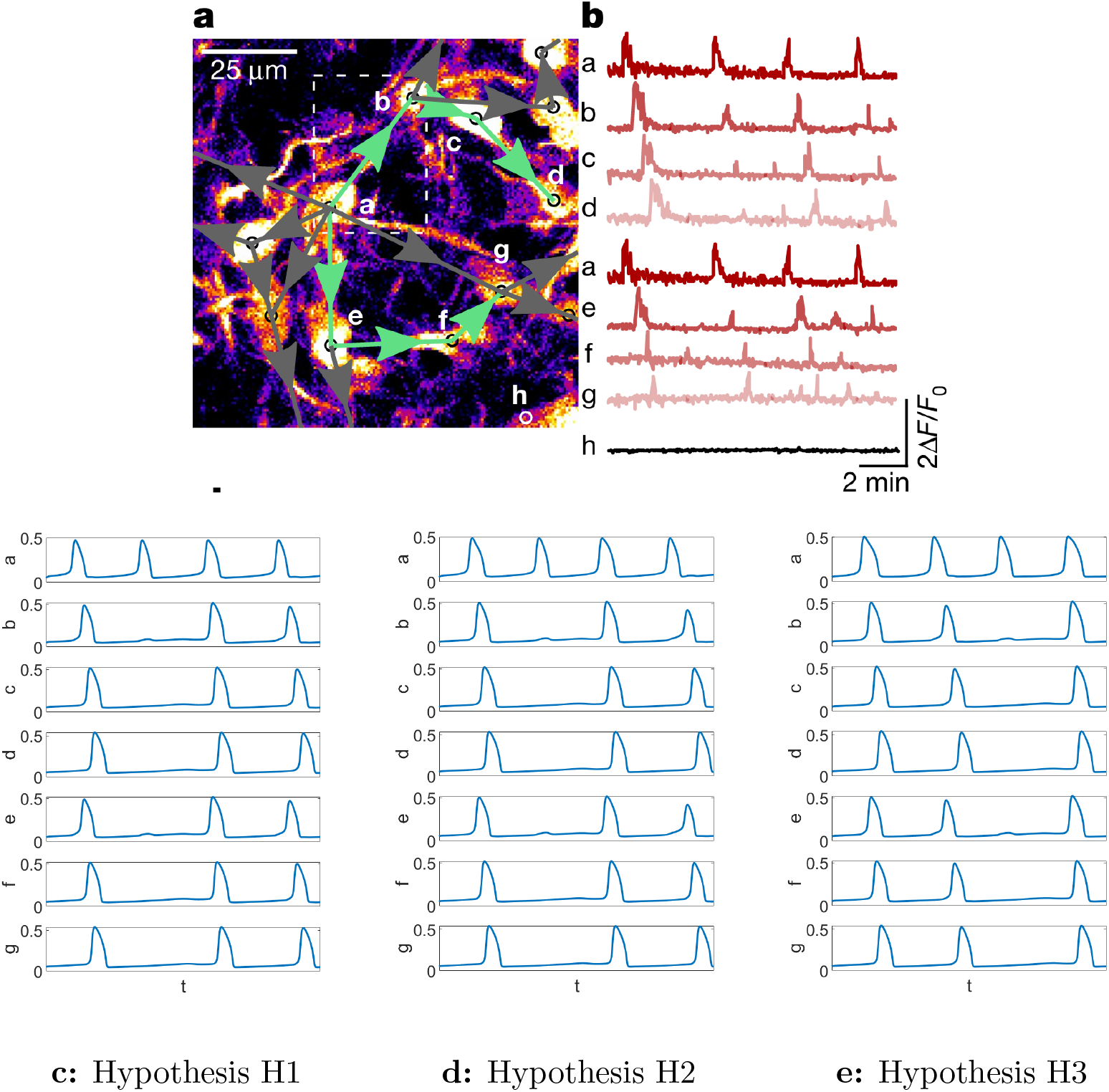
**a:** Real glioma network where cell a is periodic and the other cells are non-periodic. Cell a transmits the signal to cells b and e. Then cell b transmits the signal to cell c, which transmits to cell d. Cell e excites cell f and then f excites g. **b:** Measured calcium transients from the network. The top images **a** and **b** were taken from [17] with permission. The panels **c, d, e** show the solutions of (17) under each hypothesis H1, H2, H3 with parameters taken from Table 1. In each image, the rows illustrate the cytosolic calcium concentration of a given cell. The total time interval is 760 seconds or 12.67 minutes for each simulated row.

Note, that H2 does not account for calcium’s ability to trigger KCa3.1 pumps and hence causes an inflow of calcium through the Orai pumps. We do not account for this for simplicity.

### 4.3 Simulation of the model

In Figure 5, we show the resulting solutions of the glioma-network model (17) for each of the particular hypotheses. The network in this first example is a simple connected network comprised of three cells, where cell 1 signals to cell 2 and cell 2 signals to cell 3. Note that there are only two TMs in the network. Further, cell 1 is set to be periodic and the other two cells are non-periodic. In the first row of Figure 5, we show the resulting *J*_*in*_ for periodic and non-periodic cells based on the three hypotheses. In the second row of Figure 5 we show the solutions where the cytosolic calcium concentration with respect to time for the three cells is illustrated. We see that all three hypotheses (H1)-(H3) yield similar spike patterns where the non-periodic cells spike after the periodic cell. Further, cell 3 spikes after cell 2 as cell 2 signals to it. The spike intensities vary for non-periodic cells over time whereas the spike intensities remain fairly uniform for periodic cells for all 3 hypotheses. Further, it is possible for cell 3 to not spike at all under each hypothesis, since cell 2 sometimes produces a small calcium spike.

In Figure 6 on the top left we show a section of a real glioma network as identified in [17]. Only cell **a** is periodic. Figure 6 **b** shows the measured calcium transients of the seven cells of this network as functions of time. We can clearly see how the transients move from cell to cell, and we also notice some missed beats, as the stimulation is sometimes not strong enough to trigger a full response. We use our model (17) to test the three hypotheses H1-H3 on this network. In Figure 6 panels **c, d, e** we see that all the three hypotheses yield similar behaviors that resemble the data in the top right image. The second peak in hypothesis H1 and H2 is very small for cells **b** and **e**, and hence those cells are unable to transmit the signal further for that time interval. In contrast, hypothesis H3 shows a missing beat at the third transient.

## 5 Networks

In [17] it was shown that glioma cells in vivo have small-world and scale free network properties. It was also noted in [17] that glioma networks can have several “hub cells” which are cells with high degree (many connections) as compared to non-hub cells [30]. Networks hubs are prominent in scale-free networks and can also arise in small-world and random networks.

### 5.1 Small-world network

The small-world network is a network characterized by nodes having short paths between each other and a high clustering coefficient [30, 40]. A high clustering coefficient indicates that the nodes neighbours are likely to be connected to each other, whereas a low clustering coefficient means that the nodes neighbours are unlikely to be connected to each other [30]. Examples of small-world networks are neural network of the worm Caenorhabditis elegans and power grid of the western United States [40].

The small world network can be generated from a regular graph [30, 40]. A regular network or graph, is a graph where all nodes have equal degree and are connected to their nearest neighbours (see Figure 7). Then the small-world network is generated by rewiring each edge to a different node with probability *β*. That is, each edge in the regular graph has a small probability of *β* to be attached to a different node, while staying attached to one node in its original link. Note that 0 *< β <* 1 and as *β* → 1 the network becomes progressively more random [40].

**Figure 7:**
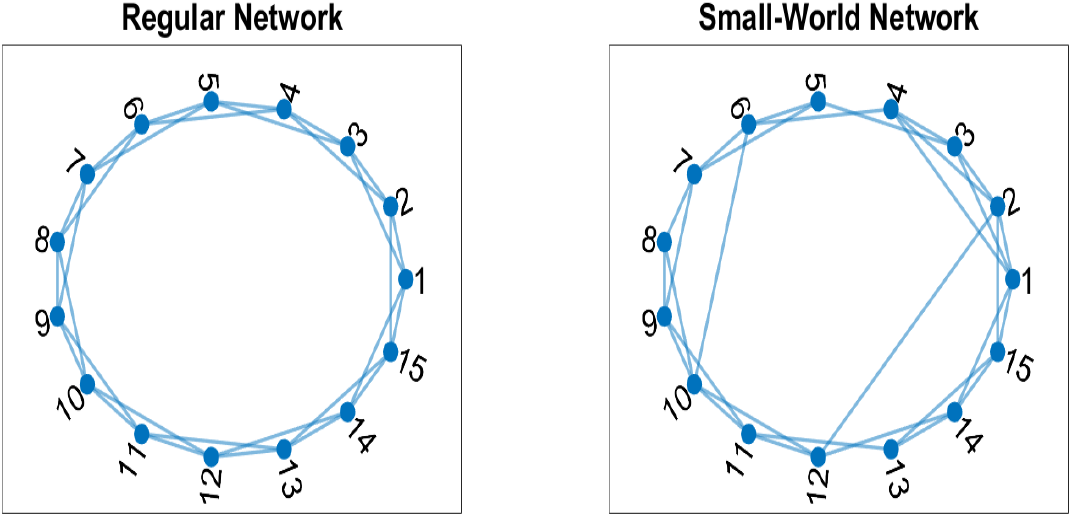
The first plot is a regular graph with degree of 4 and the second image is a small-world network with *β* = 0.2.

### 5.2 Scale-free network

A scale-free network is characterized by its nodes having a power law degree distribution, that is many nodes have low degree and few nodes have very high degree [30]. Scale-free networks also have the small world property where paths from one to another are short and the clustering coefficient is high [30]. Example of this type of network is the World Wide Web [30]. The scale free network can be generated using the Barabási–Albert (B-A) algorithm [1]. This algorithm works on the principle that nodes with higher degree or links are more likely to get chosen. Mathematically, the probability, *P*, of choosing a node, *n*, depends on its degree, *d*_*n*_, and total degree of the network, *d*_*T*_, that is

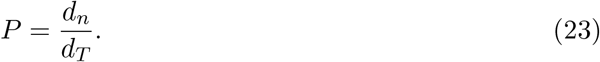

The total degree of the network, *d*_*T*_ can be calculated as

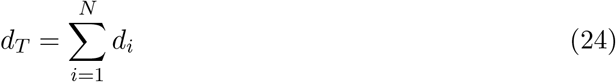

where *d*_*i*_ is the degree of node *i* and *N* is the total amount of nodes.

To generate the network, we can start with two connected nodes. Then with each new time step a new node gets added with one edge. The edge gets connected to an existing node according to the probability *P*. This generates a tree like network seen in Figure 8 (left image). Alternatively, a node with two new edges can be added to the network. In this case, the network is started with three connected nodes. The new nodes edges get attached to previous nodes according to the probability *P*, and the new edges cannot connect to the same node. This case generates a network seen in Figure 8 (right image).

**Figure 8:**
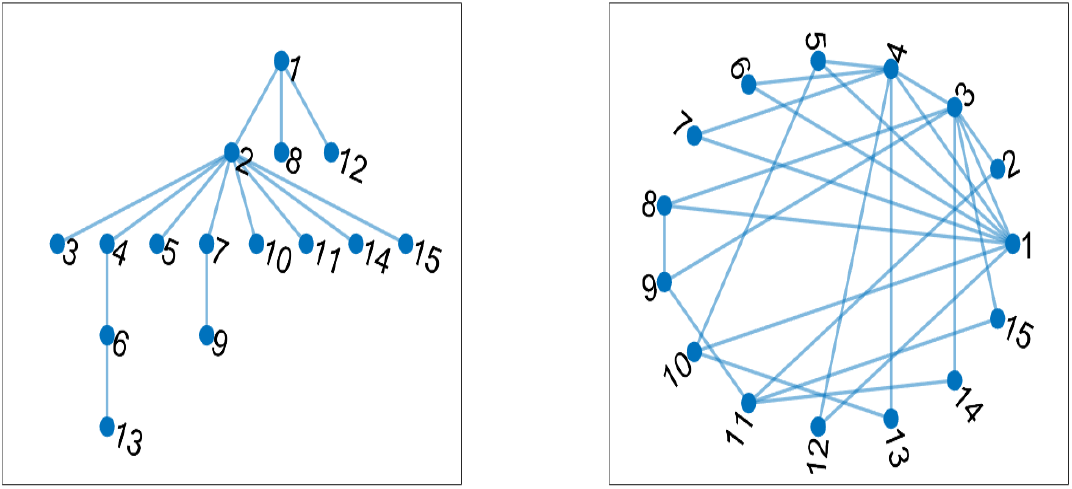
The first plot on the left denotes a scale-free network generated from adding a node with one edge iteratively, while the second plot on the right denotes a scale-free network generated from adding a node with two edges iteratively.

### 5.3 Random network

We will compare the calcium communication properties of small world and scale free networks with random networks. To construct a random digraph, cells were randomly linked to each other until the specified amount of edges was reached. Self loops and unconnected cells were removed from the network, meaning that the final amount of cells and edges differs from the initial specified amount. Edges were then added randomly to get the desired edge count. The network was then created using Matlab’s digraph function, which gives the adjacency matrix used in the model.

### 5.4 Network creation

To create the networks we use for model simulation, we follow the following procedure:

1. Create an undirected graph using the algorithms outlined above.
2. Take the upper triangular matrix from the resulting adjacency matrix from the previous step. Then randomize the directions in the digraph.

Note that taking the upper triangular matrix of an undirected graph creates a digraph with the same structure. We randomize the directions in the digraph in step 2 above to avoid predictable and biased outflow from the first nodes to the later nodes. To randomize the directions, we iteratively go through each edge and randomly switch the direction of each edge. Note that since a small-world algorithm creates more edges than a scale-free algorithm, we add extra edges randomly in between steps 1 and 2 when creating a scale-free network. This is done in order to have a consistent edge count when comparing together all networks. Further when comparing the networks in a later section, the small-world network is constructed from a regular network with each node having a degree of 4 and the scale-free network is constructed from 3 connected nodes (node 1 is connected to node 2 and node 2 is connected to node 3) and adding a node with two edges at each iteration. The two edges connect randomly to other cells and cannot connect to the same cell.

## 6 Treatments

In this section, we explore some of the treatments that were tested in [17] numerically on the various glioma networks. We test the following treatments:

T1. Random removal of cells,

T2. Random removal of TMs,

T3. Targeted removal of periodic cells,

T4. KCa3.1 pump inhibition treatment.

Of the above treatments, T1, T3, and T4 were performed experimentally in [17]. We added treatment T2, as tumor microtubes might be an alternative treatment target.

Note that when we discuss networks resistance to damage we are focusing on the networks capability to distribute the signal to other cells by measuring how many active cells there are. We define active cells as those cells that show a calcium transient once stimulated from a neighboring cell (or being a periodic cell).

### 6.1 Random treatments

We first study how removal of random cells and TMs affects the network. To do this, we work with hypothesis H2 of the model. We construct the 3 networks: small-world, scale-free, and random, all with 100 cells and 200 TMs. For each network, we first choose the networks to have 1 periodic cell and then set the networks to have 3 periodic cells. The periodic cells are chosen randomly using a random number generator. If the network has 1 periodic cell, the periodic cell is cell number 90. If the networks have 3 periodic cells, the periodic cells are cell numbers 90, 27, 87.

Figure 9 shows the resulting trends when a certain number of cells or TMs are randomly removed for each network. Rows 2 and 3 of Figure 9 show the amount of active cells after a certain percentage of cells are removed from a network with 1 periodic cell (row 2) or 3 periodic cells (row 3). We see that all three networks produce similar trends and more periodic cells do not make a significant difference in making the network more robust to random cell damage. The trend is linear, showing that the network is able to function and spread signal to all of the remaining cells, granted that the periodic cell has not been eliminated. After about 60% of cells are removed it becomes likely to remove the periodic cells and fully degrade the network. The last two rows illustrate the amount of active cells if a certain percentage of TMs are removed from a network with 1 periodic cell (row 4) or 3 periodic cells (row 5). All networks in this case yield similar trends where removing 50% or less TMs does not significantly impact the functionality of the network and most cells remain active. Moreover, the scale-free and random networks are more resistant to random TM damage than a small-world network. Overall, all networks seem to be more vulnerable to random cell damage rather than to random TM damage. Moreover, this shows that random damage to a network is not a very effective treatment as it requires removal of nearly all cells or TMs to fully degrade the network. This confirms the experiments performed in [17] where it was shown that random damage to cells was not an effective treatment.

**Figure 9:**
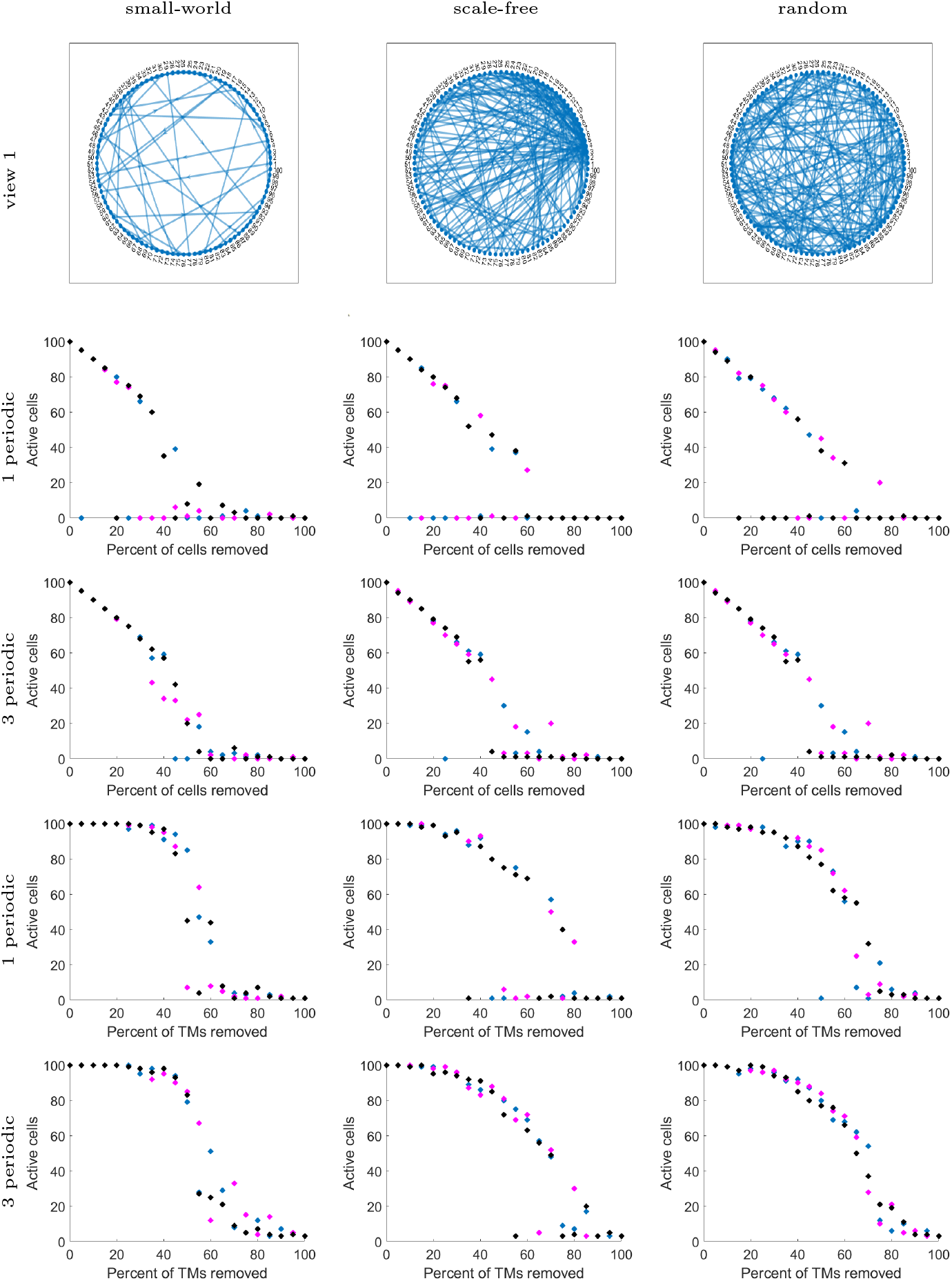
Networks with 100 cells and 200 TMs. The model used in simulations is (17) under hypothesis H2 with parameters taken from Table 1. The first row illustrates the networks. The second and third row show the amount of active cells when a random percent of cells are removed from each network. Second row uses networks with 1 periodic cell and the 3rd row uses networks with 3 periodic cells. The 4th and 5th row the amount of active cells when random amount of TMs is removed. The 4th row uses networks with 1 periodic cell and the 5th row has uses networks with 3 periodic cells. The 3 different colors (pink, blue, black) represent the 3 different runs of random removal of cells or TMs. The horizontal axis shows the percent of cells or TMs removed and the vertical axis shows the amount of active cells.

### 6.2 Targeted cell ablation treatment

In this section, we explore the targeted cell ablation where we focus on removing periodic cells. Figure 10 illustrates the amount of active cells for each network shown in Figure 9 with 3 periodic cells initially. We see that for each network, all periodic cells must be removed in order to prevent calcium signalling. Since the network remains connected after removing one or two periodic cells, it is enough for one periodic cell to trigger calcium oscillations in all cells. This shows that in order to have a successful treatment for degrading the network, all periodic cells must be removed. In the experimental results in [17] it was observed that after removing all periodic cells, the network significantly reduced in the amount of cells that are able to send out transients. However, in the experimental results it was observed that a few cells can still communicate via calcium. This is perhaps due to glioma cells having the ability to have spontaneous non-periodic calcium activity which we do not account for in our model.

**Figure 10:**
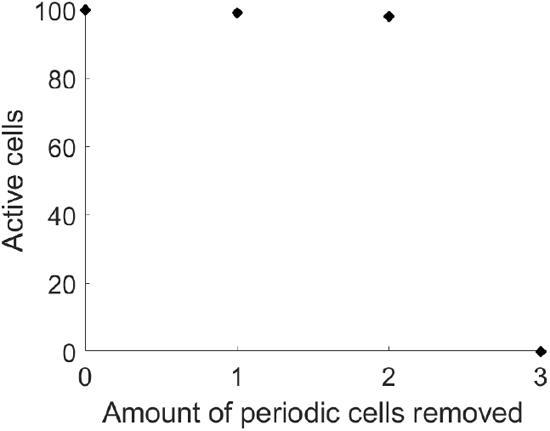
Summary of the amount of active cells post periodic cell removal for each network illustrated in Figure 9. Each network had 3 periodic cells initially.

### 6.3 KCa3.1 inhibition treatments

In Hausmann et al. [17], treatments inhibiting the KCa3.1 pump were tested. Here, we explore this treatment using the glioma-network model (17) under hypothesis H2. Under hypothesis H2, *κ* corresponds to the effect of KCa3.1 activity, therefore, inhibition of KCa3.1 treatments lower *κ*. Depending on how much *κ* is lowered, the functionality of a periodic cell and consequently the network can be significantly affected as shown in Figure 11. Biologically, the results from Figure 11 show that *κ* needs to be sufficiently lowered in order to degrade the network, otherwise the periodic cells can still signal to other cells just at lower frequency. Moreover, if the treatment inhibiting KCa3.1 has variable effect on the periodic cells, it may be possible for some periodic cells to be less affected then the others, meaning that some periodic cells signal at similar frequency as before while others signal at lower frequency or not at all. This is a potential explanation as to why the network was observed to diminish in the amount of signalling between cells in the experiments [17].

**Figure 11:**
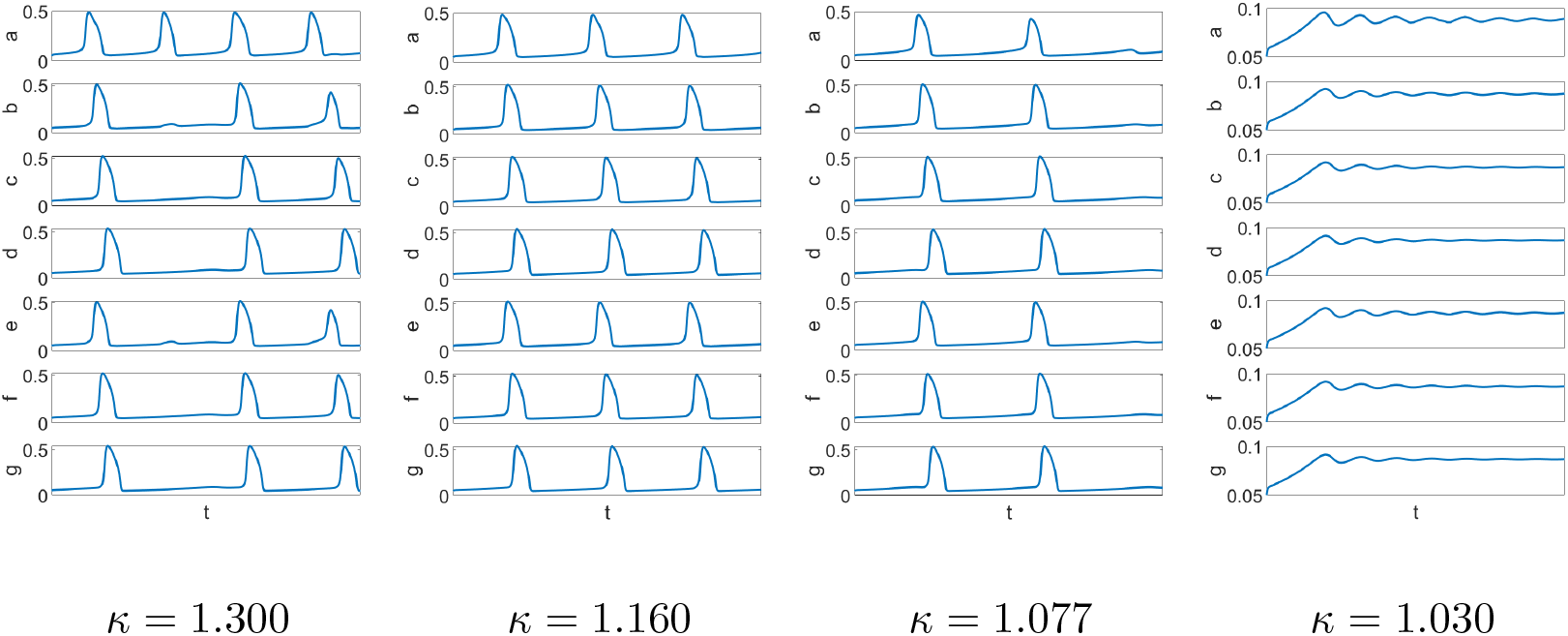
Images illustrate the respective *κ* reduction. The first image shows a simulation for the 7 cell network that was illustrated in Figure 6 under hypothesis H2 using the glioma-network model (17) (no kappa reduction). The other images show the simulation of the 7 cell network under hypothesis H2 using the glioma-network model (17) with *κ* = 1.300, 1.160, 1.077,. The total time interval is 760 seconds or 12.67 minutes for each simulated row.

## 7 Conclusion

In this work, we adapted the ordinary differential equation model proposed by Höfer [20] and accounted for two distinct cell types: periodic and non-periodic cells. We assumed that non-periodic cells cannot sustain intracellular calcium oscillations without a calcium signal from a periodic cell. We tested three hypotheses for a periodic cell having a sustained periodic intracellular calcium oscillations: a constant calcium influx into the cell proportional to the IP_3_, a constant calcium influx into the cell proportional to the IP_3_ with an added benefit from KCa3.1 pumps, or a periodic influx into the cell. In each hypothesis, the calcium oscillations stem from 3 different ways of activation. In hypothesis H1, the sustained intracellular oscillation stems from a higher IP_3_ concentration in comparison to a non-periodic cell. In hypothesis H2, the oscillations in the periodic cell are caused by a higher influx of calcium into the cell by introducing a parameter *κ* that approximates the effect of the KCa3.1 pumps. In hypothesis H3, the calcium oscillations are caused by the oscillation in the calcium influx into the cell, which is a rough approximation of the effect of the membrane potential on calcium influx. All hypotheses were able to yield similar trends in calcium oscillations. For hypothesis H3, we find that the periodic calcium influx into the cell (which was an approximation to the oscillatory calcium influx used by Catacuzzeno et al. [4]) requires a sufficiently high IP_3_ concentration (in both periodic and non periodic cells) to yield intracellular calcium oscillations. This is inline with the results of Catacuzzeno et al. [4], as they found that the IP_3_ concentration value was important for sustained calcium oscillations. We also note that in order for the model under hypothesis H2 to have sustained calcium oscillations, the IP_3_ concentration value is important. That is, small IP_3_ values require a larger *κ* value in order to yield calcium oscillations.

We showed that the glioma-network model (17) can account for occasional missing spikes in non-periodic cells. This is due to the incoming calcium signals to a receiver cell not being strong enough to trigger the calcium release from the ER. Mathematically, this is due to the perturbation from the incoming signals not being strong enough to perturb the solution trajectory away from the steady state.

We applied the glioma-network model (17) to three types of networks of different topologies: small-world, scale-free, and random. To choose the periodic cells in the network we used a random number generator. We saw through the simulations that one periodic cell is enough to sustain calcium signalling in a connected network. Interestingly, we found that if a periodic cell is a hub with a high degree, then it may not have sustained calcium spikes simply because too much calcium is flowing out of it. Therefore, periodic hub cells might not be able to cause oscillations in the network. In order to obtain sustained calcium oscillations, the parameter *γ* has to be increased.

Using the three networks we tested three different treatments: T1 random cell removal, T2 random TM removal, T3 targeted removal of periodic cells, and T4 KCa3.1 inhibition treatment. We found that the network structure has little effect on random cell or TM removal treatments, where all networks were more resistant to random TM removal than to cell removal. This is because the TM treatment is unable to remove periodic cells and it is difficult to remove all TM connections such that the periodic cells become isolated or to isolate the network hub cells. Thus, it is better to focus on a treatment specifically targeting the periodic cells, such as the KCa3.1 inhibition treatment. Using hypothesis H2, we tested the KCa3.1 treatment by lowering *κ*, which showed a decrease in calcium spike frequency, which consequently lowers the overall calcium transient frequency in the cell network. Moreover, if *κ* is sufficiently low, then the periodic cells are unable to produce calcium transients. This is in line with experimental results where they find that periodic activity decreases post a KCa3.1 treatment [17].

We note that we did not test all the treatments performed by Hausmann et al. [17]. For example, Hausmann et al. [17] studied a gap junction inhibition treatment. In the model, this treatment can be tied to random removal of TMs that we performed and to lowering of *γ* in the model. The parameter *γ* is related to the rate of transmission of calcium between cells. Making it sufficiently low will prevent the flow of calcium between cells and hence not trigger calcium oscillations in the non-periodic cells, which is a potential way of incorporating it into the model. For this study, we chose to keep all *γ* the same between all cells for simplicity. Calcium chelation was also performed experimentally in [17] outside the cell. We did not study this treatment as there is ambiguity into how it could affect the calcium influx with the hypotheses explored here. Further, the main outcome of calcium chelation was the reduction in the cancer proliferation rate which could not be tested on short time scale we studied. Finally, to study the ATP inhibition treatment, an extension to the model must be made that incorporates the ATP dynamics.

There are several limitations in the model. We did not account for the fact that IP_3_ can be transmitted through gap junctions and chose to keep the IP_3_ concentration constant in all cells (only varying the value based on the hypothesis). Further, the experimental data shows that receiver cells can have amplitudes less than the signalling cell which is a trend that is not captured well by the model. Further, there are cases in the data where the non-periodic cells do not receive a signal for a prolonged period of time after showing activity or non-periodic cells have less calcium oscillations the further away they are from a periodic cell. Based on our chosen model parameters, this behaviour is unable to be captured. However, this could be accounted for by varying the rate at which calcium gets transferred from one cell to another (*γ*) which could also vary in time. We additionally note that the parameters used are mainly taken from Höfer and their model pertained to hepatocytes. True parameters pertaining to glioma may differ and require more experimental measurements to be fully determined.

Because we studied the model on a short time interval (760 seconds) the model does not incorporate calcium’s influence on cell proliferation or take into account cell population changes over time. Further, network repair (either production of new TMs or experimental reintroduction of periodic cells) was not studied here, and hence is an outlet for future work. We also note that a more sophisticated model can potentially be created to fully characterize calcium entry into the cell, taking into account cell membrane potential as well as the various calcium channels (aside from the Orai channels).

Finally, it is still unclear whether periodic cells have qualities of cancer stem cells (which have been previously mathematically modelled [12, 18, 23, 37]) and it would be interesting to see new biological experiments that analyze the periodic cells further as well as mathematically model the interplay between periodic cells, cancer stem cells, and differentiated glioma cells.

## Acknowledgments

AS acknowledges the funding from the Alberta Graduate Excellence Scholarship and The Maud Menten Institute Accelerator award (PRN2). TH is supported through a discovery grant of the Natural Science and Engineering Research Council of Canada (NSERC), RGPIN-2023-04269.

## A Appendix

### A.1 Simplification of the Höfer model

In Section 2 the Höfer model (1) was introduced, which was then simplified to model (2). Here we show how the model can be simplified using the approach discussed in [22]. We show how to simplify the first two equations of (1) in the first equation of (2) as was done in [20]. Simplifying the last two equations in (1) to obtain the second equation in (2) is analagous.

The bound buffer is assumed to be in quasi-steady state, therefore

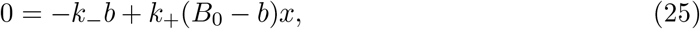

which can be solved for *b* yielding

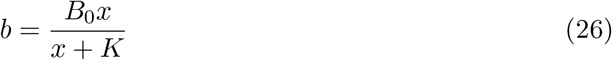

where *K* = *k*_*−*_*/k*_+_.

Adding the first two equations in (1) yields

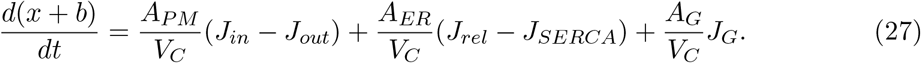

Then using (26) we rewrite (27) as

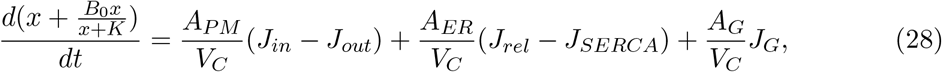

which equals to

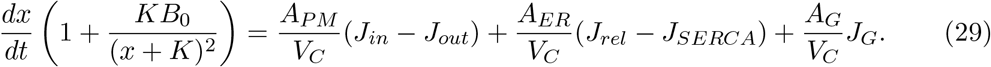

by differentiating the left hand side with respect to *x*. Simplifying (29) yields

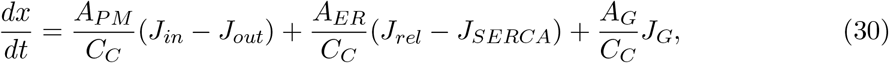

where

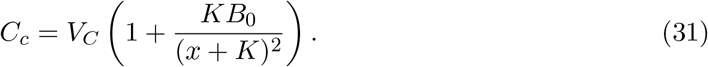

Finally, only the unsaturated case is considered in [20] meaning that *x << K*, so

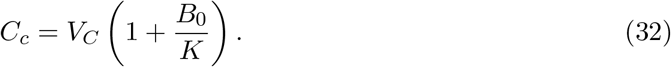

### A.2 Mathematical analysis of Höfer model

To begin, we first ignore the flux coming from external cells, hence *γ* = 0 or *J*_*G*_ = 0 and study the dynamics of a single cell. We recall the detailed model here for convenience

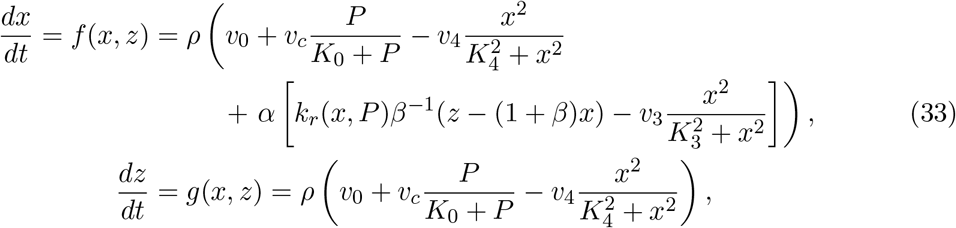

where

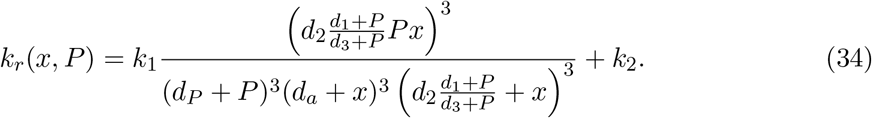

The equilibria can be determined by the intersection of the nullclines, *f* (*x, z*) = 0 (*x*-nullcline) and *g*(*x, z*) = 0 (*z*-nullcline). The *x*-nullcline and *z*-nullcline given by [20]

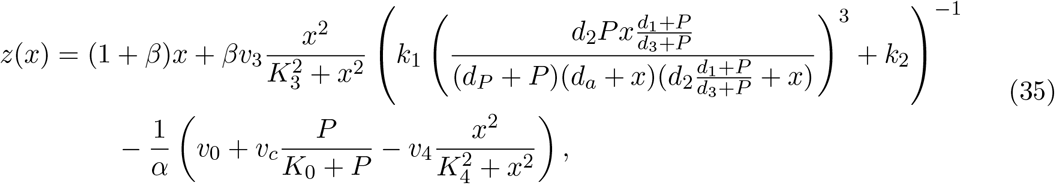

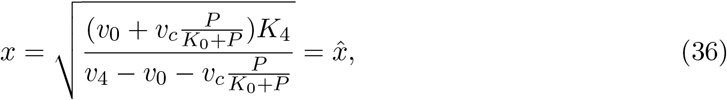

where the *x*-nullcline has been solved for *z* and the *z*-nullcline has been solved for *x*. Note that we are interested in biologically relevant equilibria, hence either 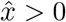 and 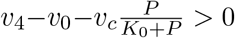 holds or 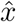 is not in the relevant domain.

Since the *x*-nullcline is a function of *x* and the *z*-nullcline is simply 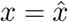 there is a unique intersection between these nullclines yielding a singular equilibrium 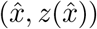. To determine its stability, we can examine the Jacobian for the system (33) which is given by

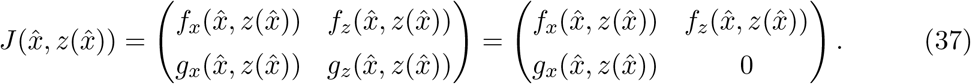

Computing the *f*_*z*_ and *g*_*x*_ we obtain that

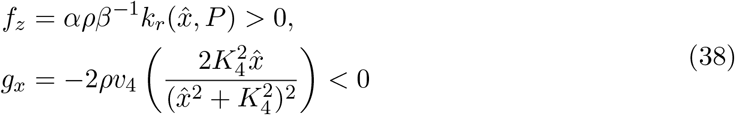

Since 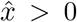. Then 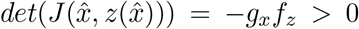 and 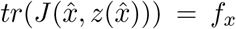. Hence, when 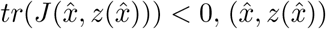 is locally stable and when 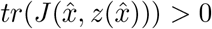 it is locally unstable. Höfer [20] showed numerically that the system (33) has a Hopf bifurcation.

In order to have a Hopf bifurcation, consider a bifurcation parameter 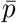. If 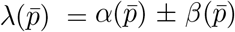 are the pair of eigenvalues for a two dimensional ODE system and there exists a parameter value 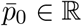 such that *β* is imaginary, 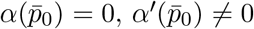 and 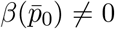 then 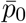 is a Hopf bifucation point [22]. The periodic solutions resulting from the bifurcation are stable if in addition 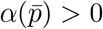.

From (37), we find that

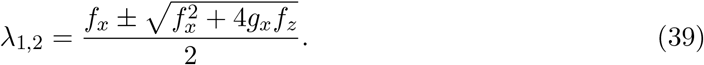

We set

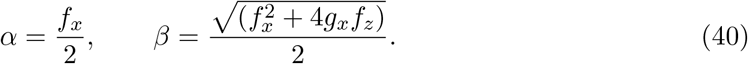

Notice that if *β*^2^ *<* 0 then we have a pair of complex eigenvalues and if *β*^2^ *>* 0 then we have a source or sink node (saddle is impossible since 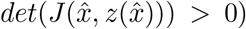. We use IP_3_ concentration *P* as a bifurcation parameter as done in Höfer [20]. We plot *α*(*P*), *β*^2^(*P*), and *α*^*′*^(*P*) in Figure 12 using parameters from Table 1 while varying *P*. Höfer stated that *P* = 1.45 and *P* = 8.892 yield a supercritical Hopf bifurcations (stable periodic solutions) which is confirmed in Figure 12 since at those points *α*(*P*) = 0, *β*(*P*)^2^ *<* 0, *β*(*P*) ≠ 0, and *α*(*P*) *>* 0 when *P* ∈ (1.45, 8.892). However, we notice that in the interval *P* ∈ (1.45, 8.892), *β*^2^(*P*) becomes positive when *P* ∈ (2.594, 5.33) meaning that the equilibrium becomes a source instead of a spiral. Numerically, we still get stable periodic orbits in that region as shown in Figure 3 for *P* = 3 and *P* = 5. Figure 3 illustrates some example phase portraits to supplement the analysis, where we see that trajectories tend to follow the *x* nullcline closely before converging to the equilibrium or approach a periodic orbit. We also note that *P* is not the only bifurcation parameter due to the complexity of *f*_*x*_ and there are other possible parameters to yield a Hopf bifurcation.

**Figure 12:**
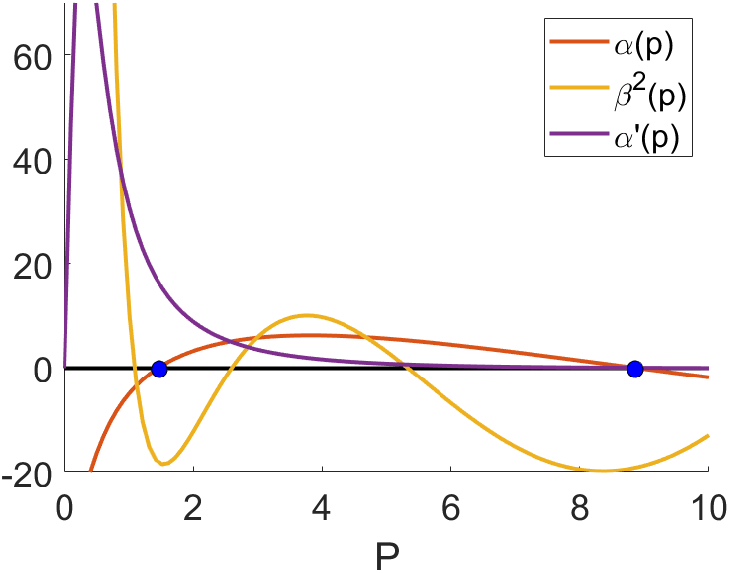
Illustration of *α*(*P*), *β*^2^(*P*), *α*^*′*^(*P*) using parameters from Table 1 where *P* varies. The blue dots indicate the Hopf bifurcation points.

From Figure 3, we notice that the model results in an excitable system when the steady state 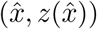 is stable, that is *f*_*x*_ *<* 0. An excitable system, by definition, is a system which has a stable steady state and a perturbation away from the equilibrium can send the solution orbit on a long excursion [22]. An example of a classic excitable system is the FitzHugh–Nagumo model [22], which also has a cubic nullcline for the first compartment (the nullcline which solution trajectories follow) and a linear nullcline for the second compartment. Depending on the parameters, the FitzHugh–Nagumo model can yield solutions converging to a steady state (and when perturbed the solutions can go on a long excursion) or to a stable orbit.

Based on the phase portraits, we notice an issue with the model that Höfer [20] did not discuss, which is the possibility for the solutions to escape the positive quadrant through the *x*-axis (*z* = 0) yielding solutions with negative concentrations. This is evident from Figure 13 which shows a zoom into the positive quadrant. Before the *z* nullcline (illustrated in pink in Figure 13) the vectors always point upward but beyond the *z* nullcline the vectors always point downward. Hence, given an initial condition with a sufficiently large *x* and a sufficiently small *z* the solution will escape the positive quadrant and turn negative before returning back to the positive quadrant to converge to a steady state or cycle. The issue is combated by picking a biologically relevant initial conditions. We recall that 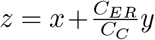 and hence *z* ≥ *x* since *y* ≥ 0. Therefore, choosing an initial condition in the grey region (*z < x*) in Figure 13 is irrelevant and hence negative concentrations do not occur for biologically realistic initial conditions.

**Figure 13:**
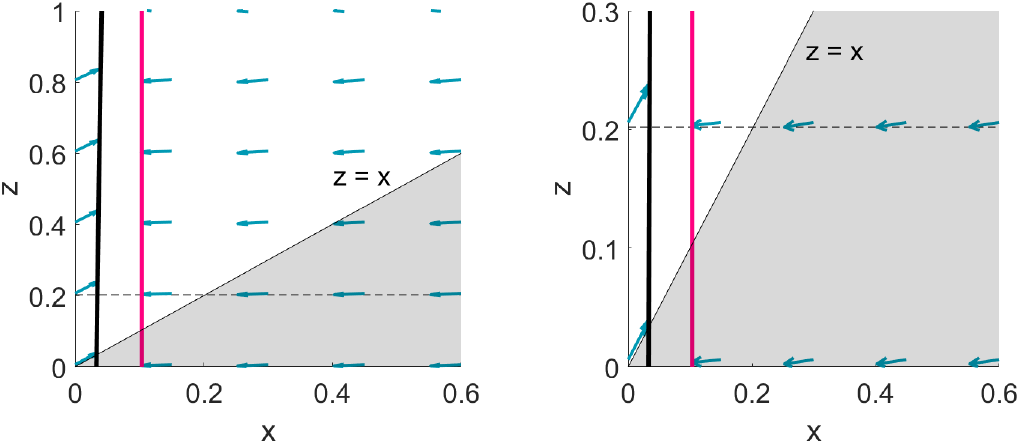
Phase portraits with parameters used from Table 1 with *P* = 2 where the right image is a zoom in of the one on the left (dashed black line is a horizontal helper line for visualization). The pink line is the *z* nullcline and the black curve is the *x* nullcline. The shaded region highlights everything below *z* = *x*.

